## Supplementary Information for "Direct observation of independently moving replisomes in *Escherichia coli*"

**This SI includes:**

Supplementary Figures 1-15.

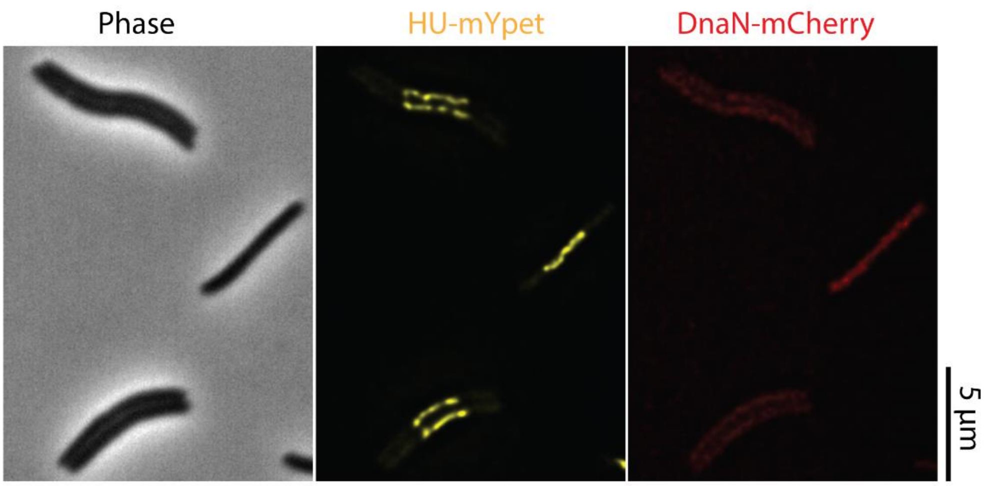


**Supplementary Fig. 1.** Temperature sensitive *E.coli* cells grown at 40ºC for 3 hours. Cells typically grow longer than rod-shaped cells while maintaining a single nucleoid that is positioned in the middle of the cell (nucleoid labelled with HU-mYpet). There are no DnaN-mcherry foci visible (only a weak background RFP signal), indicating that there are no ongoing replication rounds in these cells.

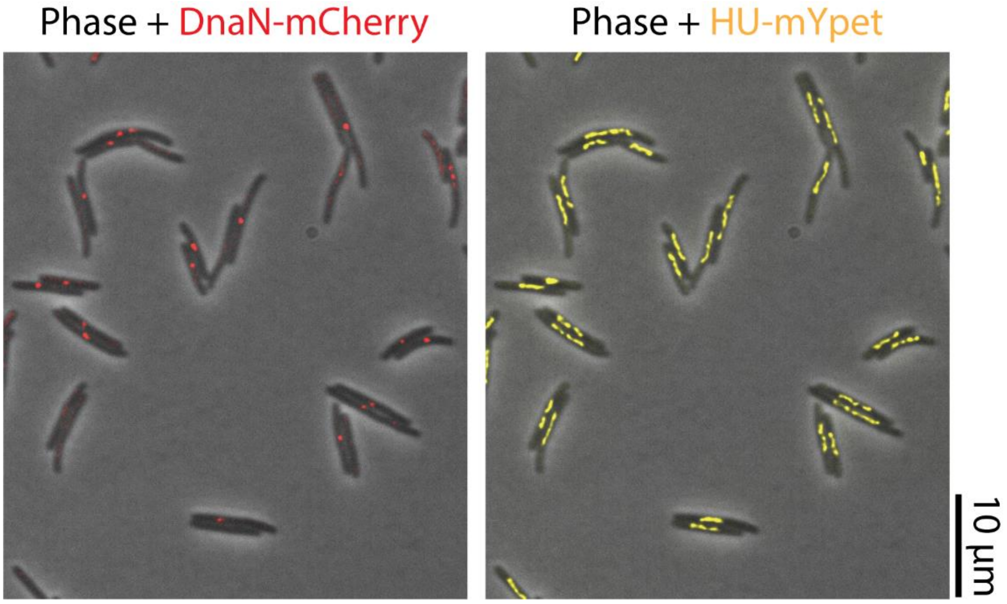


**Supplementary Fig. 2.** By placing the cells from permissive temperatures to room temperature for 10 minutes, one can synchronously re-initiate replication in ~85% of cells evidenced by the formation of replisome foci (red spots) in the DnaN-mcherry channel (left).

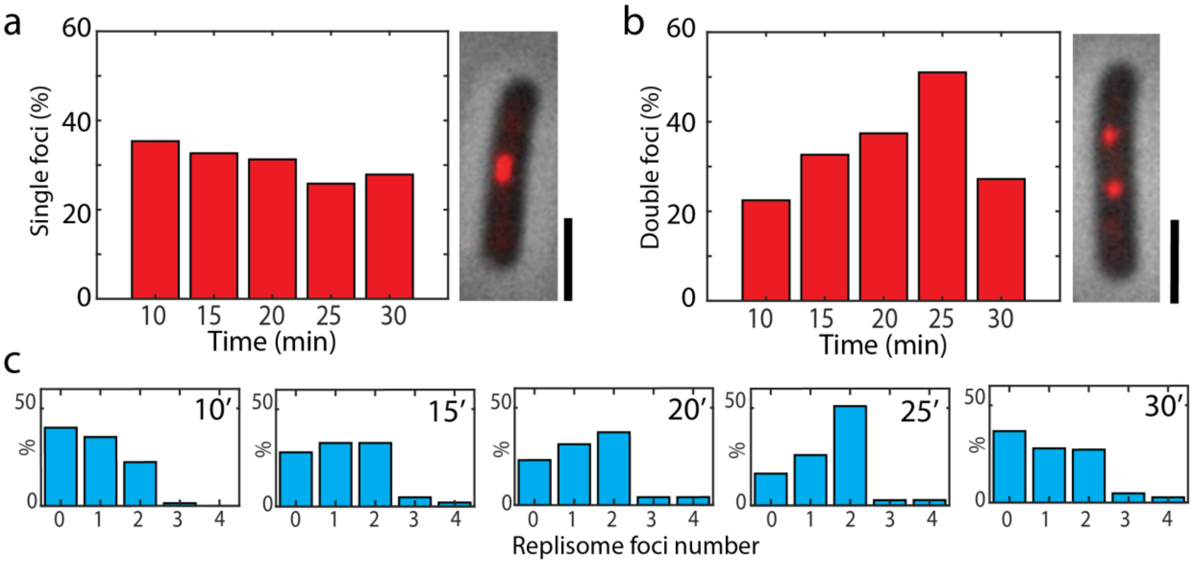


**Supplementary Fig.3 a.** The number of cells having single or **b.** double replisome foci during the first 30min of replication. While the number of cells with single foci is not changing significantly, the number of cells with two foci is gradually increasing, reaching a maximum (~50% of all cells) at time point T=25 minutes after replication initiation (*N=167*). Inset shows overlay of images of *E.coli* cells in the phase and DnaN-mCherry channels. Scale bars 5μm. **c**. Number of all replisome foci vs the imaging time. Time indicated in minutes.

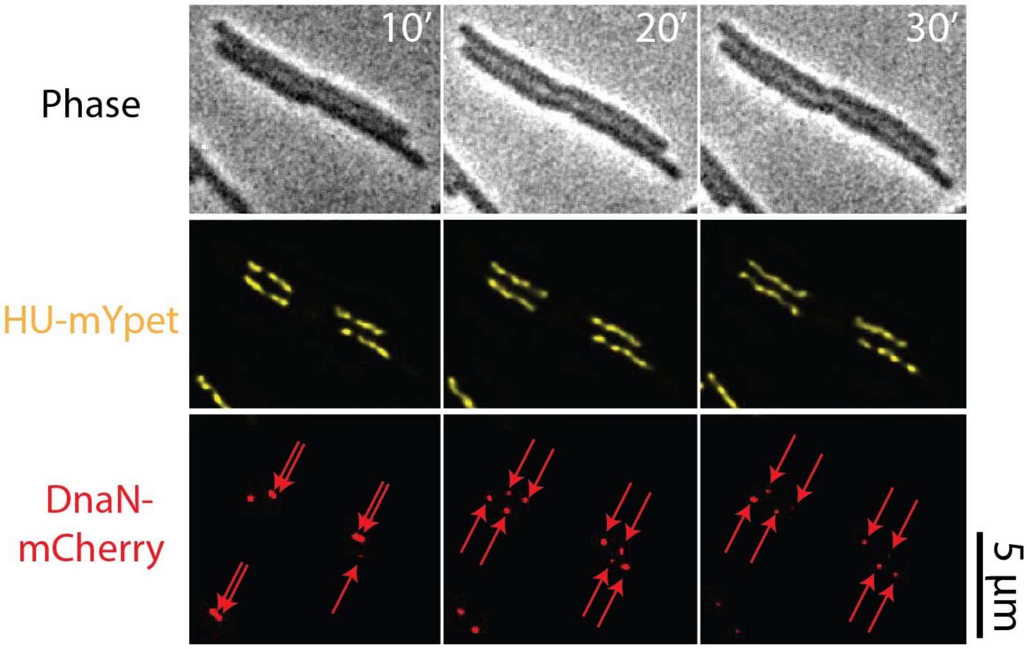


**Supplementary Fig. 4.** Time-lapse phase-contrast and fluorescent (HU-mYpet and DnaN-mCherry) images of replicating cells with two nucleoids at 40ºC. Both nucleoids replicate and the replisome foci (red spots) duplicate and move away from each other (indicated with red arrows). Time indicated in minutes.

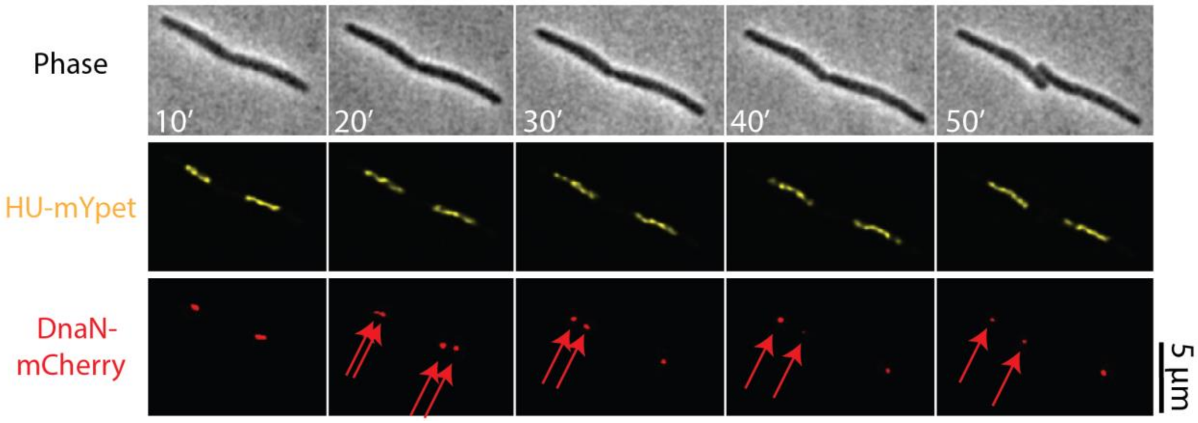


**Supplementary Fig. 5.** Time-lapse phase-contrast and fluorescent images of replicating cells grown at 37ºC. Once the cells are longer, the replisome foci (indicated with red arrows) duplicate and move away from each other. Scale bar 5μm.

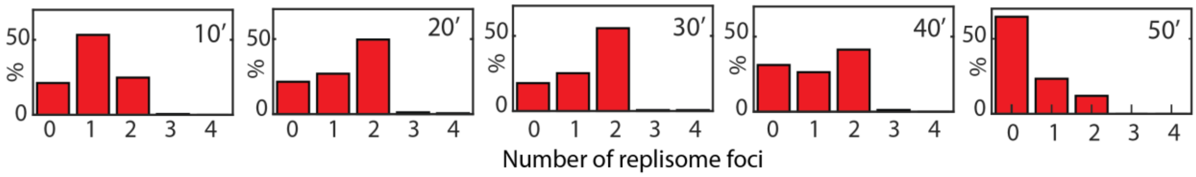


**Supplementary Fig. 6** Number of replisome foci in cells throughout the first 50 minutes of replication. A single replication focus preferentially splits in two foci throughout imaging (at time point T=30minutes, ~55% of all cells have two replisome foci).

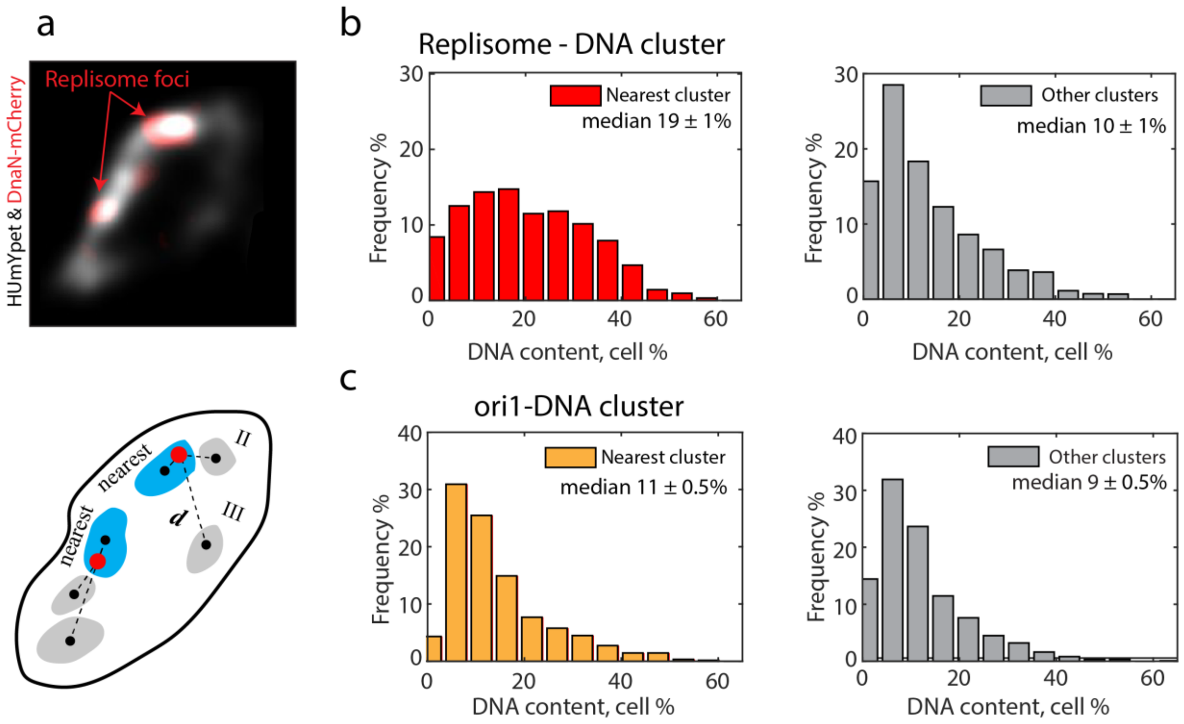


**Supplementary Fig.7 a.** Fluorescent image (top) and a schematics (bottom) showing the position of replisome spots (red circles) and DNA clusters (coloured areas) inside a cell (cell contour drawn as continuous black contour)[1]. Dashed lines indicate the respective distances *d* between the replisomes and the nearest (cyan) or any other (grey) clusters. **b.** Replisomes are found to be preferentially positioned near the largest DNA clusters (N=257 cells analysed). The relative DNA content in the cluster nearest to the replisome foci typically contains ~19% of the total DNA signal (red bars), compared to only a ~10% size for random clusters (grey bars). The values are median values ± s.e.m. **c.** If, as a control, we instead track the ori1 positions rather than the origin of replication, the preferential clustering is lost (orange bars) (N=561 cells analysed), as both the nearest (orange bars), as well as any other clusters (grey bars), contained roughly ~10% of the total DNA.

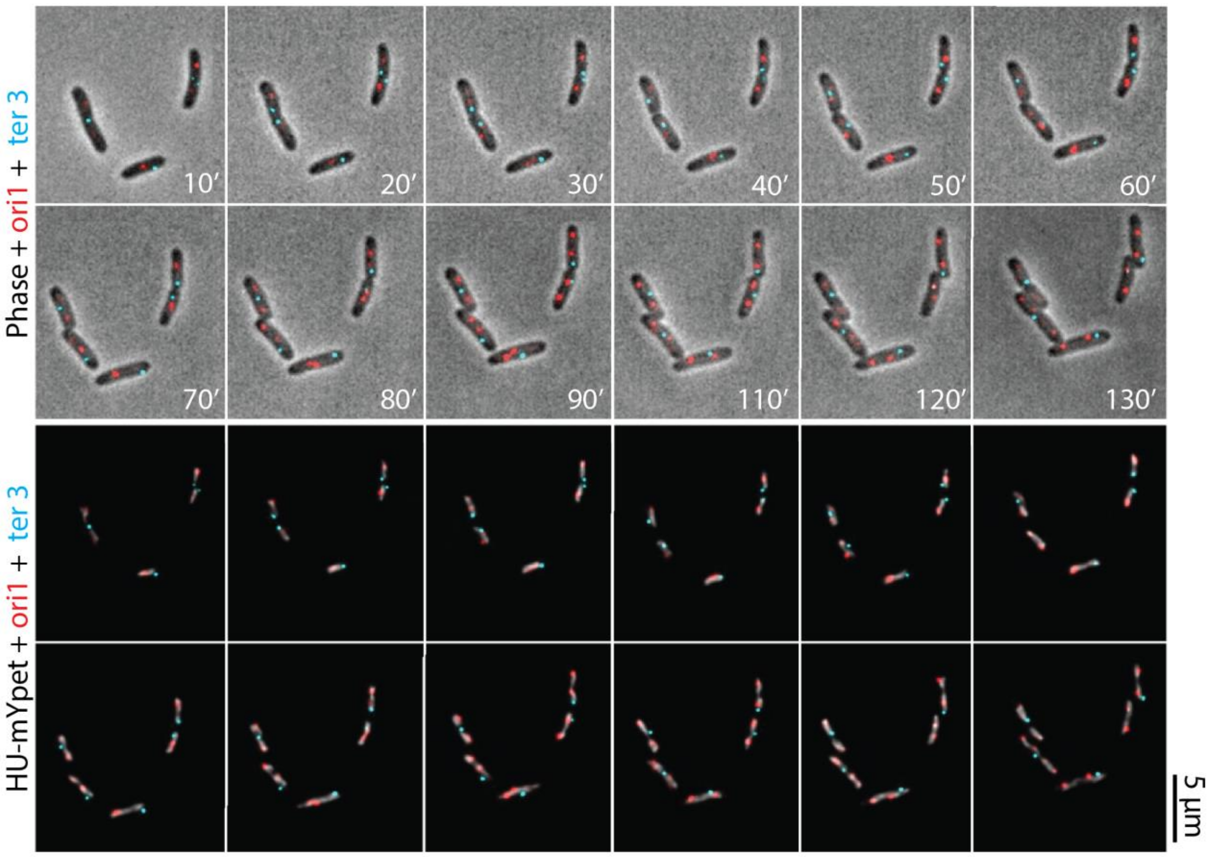


**Supplementary Fig. 8** Time-lapse images of replicating rod-shaped *E. coli* cells at 30ºC. Top: Overlay of phase, RFP (ori1 foci), and CFP (ter3 foci) channels. After the ori1 foci are replicated, they quickly move towards the poles. The Ori foci move passes the ter foci (cyan), which moves towards the middle of the cell, where it will stay for a longer time until it duplicates. Bottom: Overlay of YFP, RFP, and CFP fluorescent channel images. The chromosome (grey) gradually elongated as it was being replicated. At the later time points, two separate chromosomes were already distinguishable. Scale bar 5µm.Time is indicated on the images.

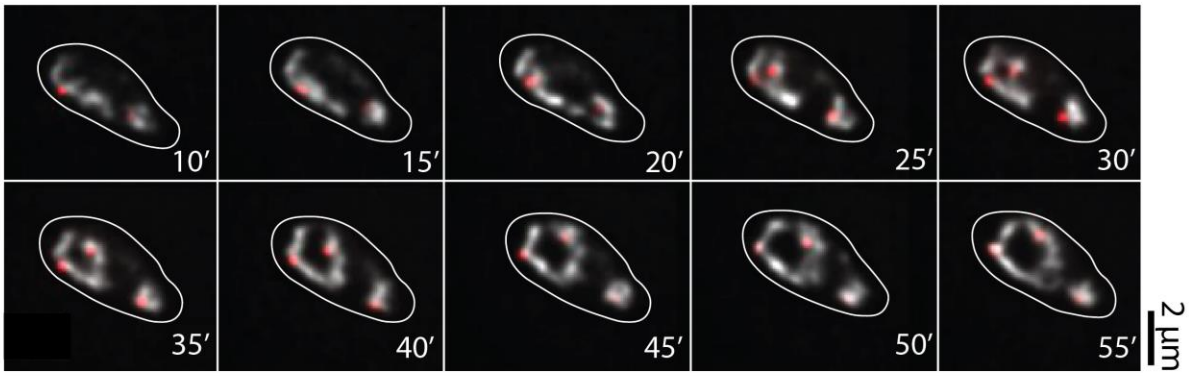


**Supplementary Fig. 9.** Time-lapse fluorescent images of a replicating cell with two nucleoids. In this example, the Ori's (red spots) did not split in the middle of the cell since their initial positions were away from the midcell. Interestingly, only the expanded nucleoid replicated and the second nucleoid stayed compacted during replication initiation. DNA was labelled with HUmYpet in grey scale and ori1 foci labelled with mCherry in red. The cell contour is shown as continuous white line and time is indicated in minutes.

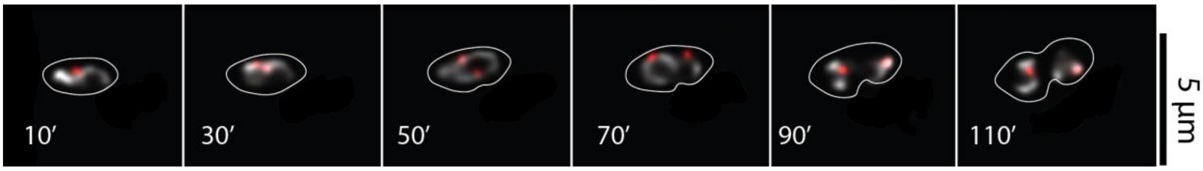


**Supplementary Fig. 10.** Time-lapse fluorescent images of replicating cell with HU-mYpet chromosome labels and ori1 foci labelled with mCherry (in red). After splitting, the Ori foci first oriented along the short axis (50' min) of the cell, and only after 70min of imaging, they re-orient towards the long axis of the cell. The cell contour is shown as continuous white line and time is indicated in minutes.

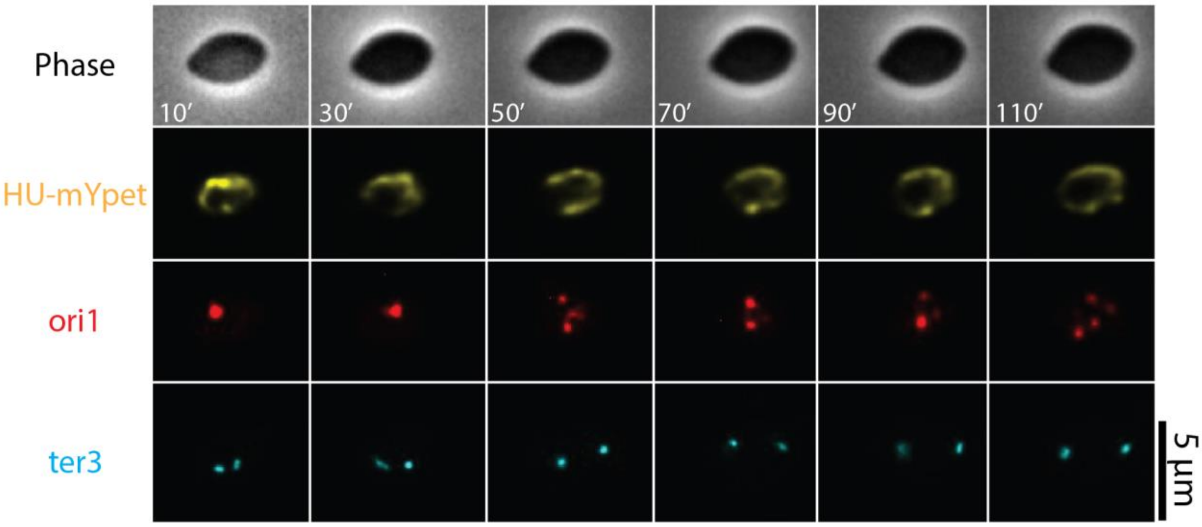


**Supplementary Fig. 11.** Time-lapse phase and fluorescent images of cell replication with segregation defects (chromosome labelled with HUmYpet, ori1 foci labelled with mCherry, ter3 foci labelled with mCerrulean). Despite Ori splitting, even after 110min the chromosomes stayed catenated, did not segregate towards cell poles and got stuck in the middle of the cell.

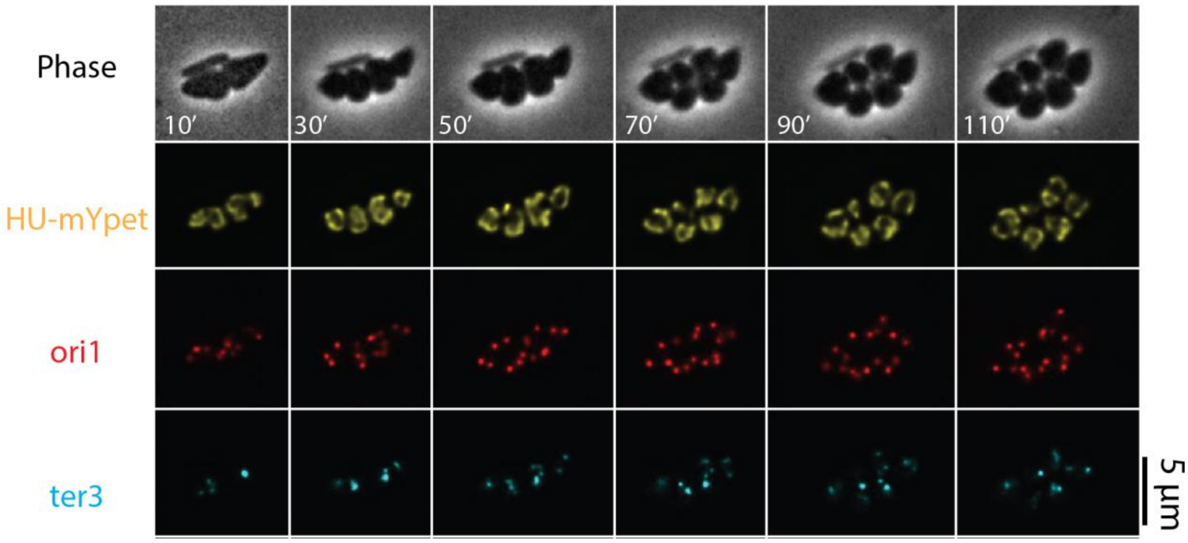


**Supplementary Fig. 12.** Time-lapse phase and fluorescent images of cell replication with segregation defects (chromosome labelled with HUmYpet, ori1 foci labelled with mCherry, ter3 foci labelled with mCerrulean). Increasing the number of replication initiations led to correct segregation in widened cells.

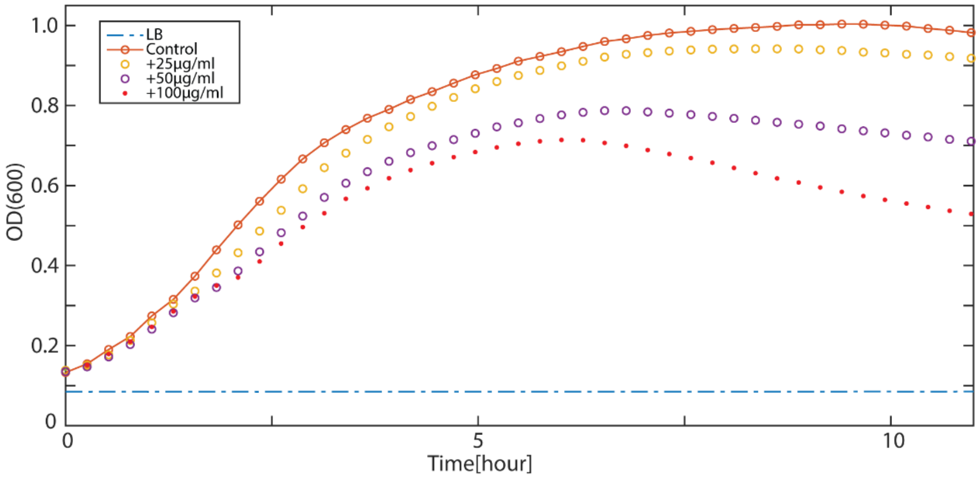


**Supplementary Fig. 13.** Cell growth curves measured by the optical density at 600nm in LB at 30ºC, in the presence of various concentrations of novobiocin drug (25µg/ml, 50µg/ml and 100µg/ml). The monitored cells and their corresponding curves are indicated in the legend. Increasing novobiocin concentration led to decreased growth rates of cells compared to control cells.

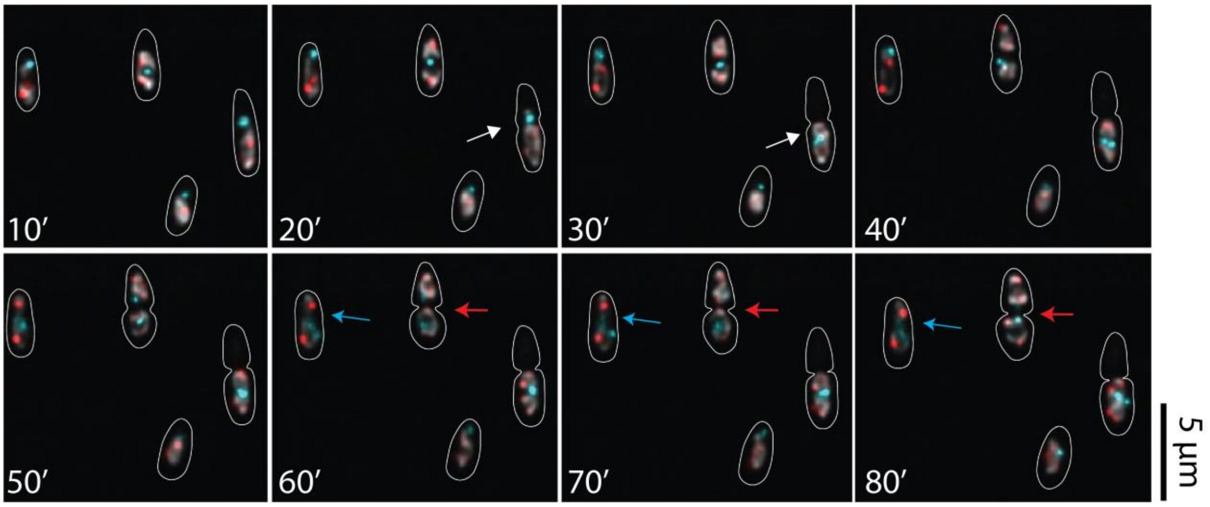


**Supplementary Fig. 14.** Time-lapse images of replicating cells in the presence of Novobiocin (50µg/ml). Overlay of HUmYpet (YFP), mCherry (RFP), and mCerrulean (CFP) channel images. The cell wall contour is indicated by a continuous white line. Adding Novobiocin had a clear influence on the replication and segregation process in cells. In the cell indicated by a white arrow, the *dif* focus (cyan foci) moved to the middle of the cell to be replicated (t=20'). During septation, both *dif* sister foci became visible in one of the daughter cells. As the result, one of the cells became anucleated. In the cell marked with blue arrow, the chromosome was positioned in the middle of the cell and the septum could not form. In the cell indicated with the red arrow, the sister chromosomes migrated to opposite cell halves, but still were connected with a thin string of un-decatenated chromosome (t=80').

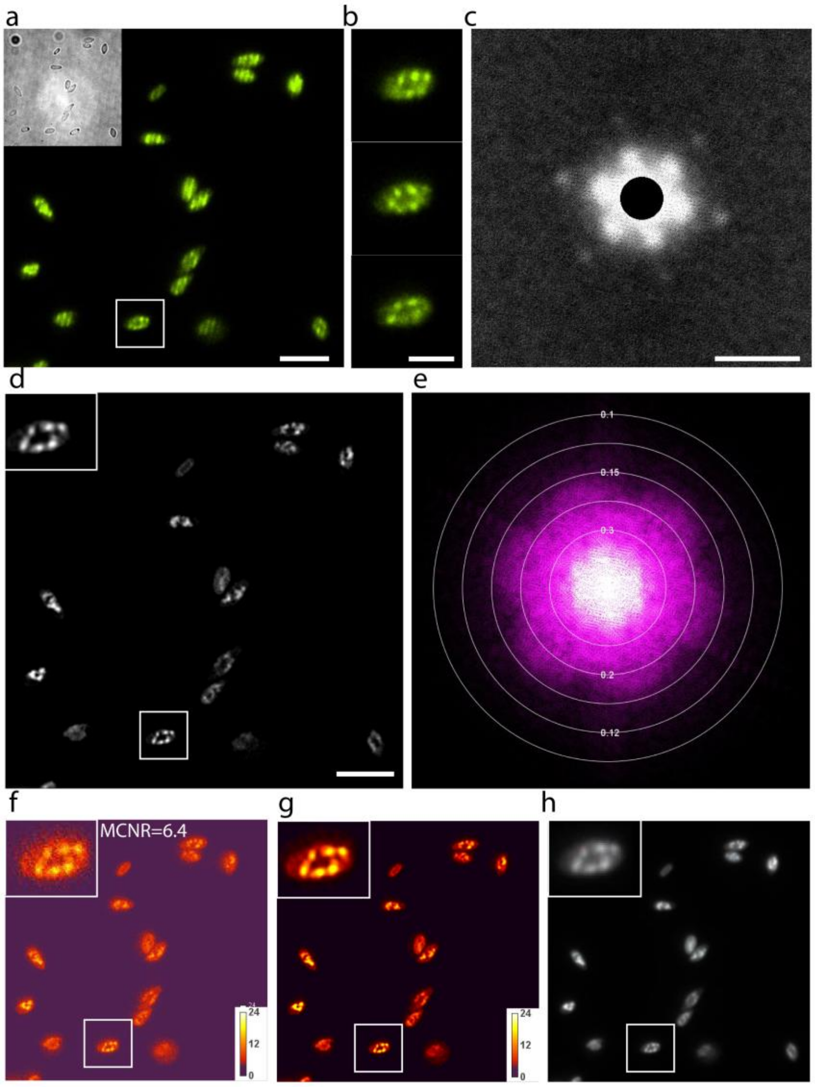


**Supplementary Fig. 15.** SIM images were tested for possible imaging and reconstruction artefacts, using the SIMcheck software[2]. **a.** Typical example of a raw SIM image of widened *E.coli* cells with HU-mYpet labelling. Inset shows a brightfield image of the same sample. Scale bars, 5 µm. **b.** Zoomed image of the cell highlighted in panel A. The dark grid-like modulation pattern in the three images shows each of the three illumination angles. Scale bar, 2 µm. **c.** Raw Fourier Projection of the raw data in reciprocal space, with points of high-frequency information from first (large inner spots) and second (outer smaller spots) order stripes. Scale bar 16 $\mu m^{-1}$. **d**. Reconstructed SIM image from the image in panel A. Inset shows a zoomed image of the circular chromosome from panel B. Scale bar, 5 µm. **e**. Reconstructed Fourier Plots overlaid with concentric rings that indicate the corresponding spatial resolution in micrometers. Based on the profiles one can approximate the effective resolution limit of features on the reconstructed data to be ~ 0.15µm. **f** & **g**. Modulation Contrast Maps for Raw (panel F) and reconstructed SIM images (panel G). The higher the value of the modulation contrast-to-the noise ratio (MCNR) (i.e., the higher the yellow-to-red color intensity) the higher the quality of SIM reconstruction. The modulation contrast-to-noise ratio (MCNR) for the full raw image was equal to 6.4, which is a satisfactory value (1), also considering that we image live bacteria and not fixed cells (1). **h**. Motion & Illumination Variation assembly of phase-averaged and intensity-normalized images for each angle in SIM microscopy. The grey-white appearance of the output image indicates motion stability and evenness of the illumination, meaning that the movement of the chromosome is slow compared to the imaging acquisition time.
